# In silico analysis of metabolic burden effects on a multicellular integral controller

**DOI:** 10.1101/2024.11.12.623154

**Authors:** Giovanni Campanile, Vittoria Martinelli, Davide Salzano, Davide Fiore, Mario di Bernardo

## Abstract

Metabolic burden is a critical limiting factor in the design of synthetic circuits, affecting both their reliability and performance. To mitigate its effects, distributing control functions across different cell populations within a multicellular control architecture offers a promising solution, while simultaneously enhancing modularity and re-usability of the circuits. We first present a model that explicitly accounts for limited ribosome availability within cells. Using this framework, we then derive a mathematical model of a multicellular antithetic integral controller that incorporates these shared resources. Through numerical bifurcation analysis and *in silico* agent-based experiments in BSim, we compare the multicellular controller against its traditional single-cell (embedded) implementation, evaluating both resource utilization and stability.

## I. Introduction

Synthetic biology aims to create novel functionalities in living organisms through engineered genetic networks. Applications span diverse fields including medicine, agriculture, energy, and environmental protection. Engineered bacteria, for instance, can detect and target cancer cells [1], [2], produce bio-fuels [3], or remediate harmful substances from water and soil [4]. However, the inherent nonlinearity, stochasticity, and variability of biological systems can limit their practical applications. Control theory offers powerful tools to address these limitations, enhancing the stability, robustness, and performance of synthetic biological systems. This interface between control and biology has given rise to *Cybergenetics*, which focuses on developing biochemical controllers for engineering reliable and robust synthetic cells. A crucial element in this field is the implementation of an integral action, which enables synthetic circuits to achieve and maintain desired outputs despite disturbances—a property known as *robust perfect adaptation (RPA)* [5].

The first Antithetic Integral Controller (AIC), introduced in [5], utilizes a pair of proteins that bind with high affinity to form an inert complex. These proteins are produced in proportion to the reference signal and the output gene requiring regulation. While both numerical and experimental studies have demonstrated this controller’s ability to achieve RPA [5], [6], implementing all control components within single cells can compromise stability and performance. A primary constraint is metabolic burden, that is, the competition between engineered networks and native cellular processes for limited resources [7]. This burden can impair cellular physiology, leading to reduced growth and increased mutation rates [8], [9], while also introducing unexpected non-regulatory interactions among chemical species [10]. Patel et al. [11] developed a model that accounts for limited ribosome availability in gene networks. When applied to the antithetic integral controller, their analysis revealed how metabolic burden and its induced non-regulatory interactions can destabilize the circuit.

A promising approach to mitigate metabolic burden is the distribution of control functions across different populations within a microbial consortium, implementing a multicellular control architecture [12]–[14]. It has been claimed that this strategy can enhance modularity and re-usability of engineered cells while enabling separation of incompatible chemical reactions and reducing the metabolic load on individual populations. Consequently, microbial consortia can achieve improved production throughput and survivability.

This paper presents an *in silico* investigation of metabolic burden effects on both embedded and multicellular implementations of the AIC. We first derive a model that explicitly incorporates intracellular resource competition in the embedded controller. We then develop a multicellular implementation comprising two microbial populations that communicate via orthogonal quorum sensing molecules. After deriving a resource-aware model of the consortium, we analyze equilibrium stability through numerical bifurcation analysis. The multicellular design is validated using *BSim*, a realistic agent-based simulator [15], with *in silico* experiments. Finally, we compare both architectures in terms of regulation capability and resource utilization efficiency.

## II. Background

This section reviews key concepts of the antithetic integral controller in two configurations: an *embedded* version, where both integral control and target processes reside within a single cell, and a *multicellular* version, where these processes are distributed across two distinct cell populations. We also present the mathematical framework used to model the effects of limited ribosome availability. Throughout this paper, we use capital letters (e.g., *X*) to denote genes and proteins, and lowercase letters (e.g., *x*) to represent their concentrations.

### A. Embedded AIC

Robust perfect adaptation (RPA)—the ability to maintain stable, predictable behavior across varying environmental conditions and cellular states—is essential for engineered biological systems. The antithetic motif [5], a synthetic gene network capable of integral control action, has been demonstrated to achieve RPA. This controller employs two high-affinity sequestering biomolecular species, *Z*_1_ and *Z*_2_, to regulate a biological process involving genes *X*_1_ and *X*_2_, where *X*_2_ production depends on *X*_1_. The feedback loop is completed by *Z*_1_ promoting *X*_1_ production and *Z*_2_ being produced proportionally to *X*_2_. Under the assumption of unlimited cellular resources, the system dynamics can be described as [16]

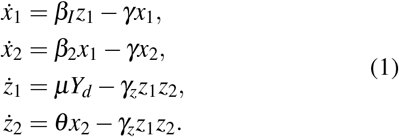

In this model, *β*_*I*_, *β*_2_, *µ* and *θ* represent the activation rates of *X*_1_, *X*_2_, *Z*_1_ and *Z*_2_ respectively, with *β*_*I*_ serving as the integral control gain. The dilution rate of *X*_1_ and *X*_2_ is denoted by *γ*, while *γ*_*z*_ represents the binding rate between *Z*_1_ and *Z*_2_, assumed to be their sole degradation mechanism, following [16]. The reference signal *Y*_*d*_ is either provided externally or encoded constitutively in the *Z*_1_ promoter. The difference (*z*_1_ − *z*_2_) is proportional to the integral of the control error *e*(*t*):= *µY*_*d*_ − *θx*_2_, thus implementing integral control [5].

### B. Multicellular AIC

A multicellular implementation of the AIC can be derived from the multicellular PI control architecture in [13] by removing the proportional controller cells. This design separates the integral control action and the biological process into two distinct populations: a *Controller* population and a *Target* population (see Fig. 1). Communication between populations occurs via two orthogonal quorum sensing molecules: *Q*_*u*_, which delivers the control input to target cells as an actuation signal, and *Q*_*x*_, which acts as a sensing signal, effectively closing the feedback loop.

**Fig. 1:**
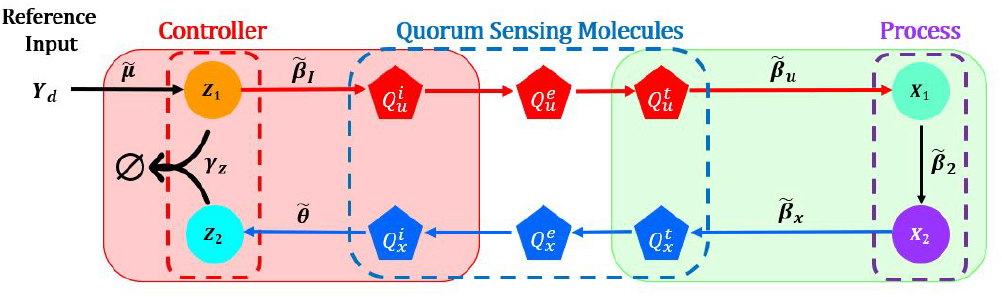
Architecture of the multicellular AIC. The genetic circuit is distributed across two microbial populations: the Controllers (in red) host species *Z*_1_ and *Z*_2_, the Targets (in green) host species *X*_1_ and *X*_2_. Orthogonal quorum sensing molecules *Q*_*u*_ and *Q*_*x*_, the red and blue polygons, respectively, implement the communication protocol by diffusing into the environment.

Throughout the rest of the paper, superscripts *e, t*, and *c* denote quantities in the environment, Target cells, and Controller cells, respectively.

The Target cells contain the same controlled process described in Section II-A, comprising genes *X*_1_ and *X*_2_. The molecule *Q*_*u*_ activates *X*_1_, while *Q*_*x*_ (the measured output) is produced in proportion to *X*_2_. Assuming the diffusion dynamics of the quorum sensing molecules to be linear [17], the dynamics of the species within the target cells can be described by

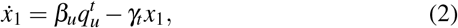

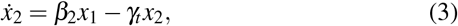

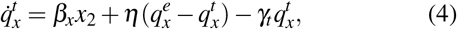

where *β*_*u*_ and *β*_*x*_ are the activation rates of *X*_1_ and *Q*_*x*_, respectively, and *γ*_*t*_ is the dilution rate of the target cells. Additionally, *η* is the diffusion rate of the molecules across the cell membrane, assumed to be the same for all cells [18].

The quorum sensing molecule *Q*_*u*_ is produced by the controller cells embedding the antithetic motif described in Section II-A. The dynamics of the network hosted within the controller cells can thus be written as

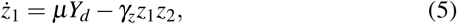

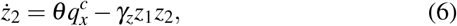

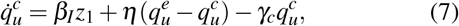

where *β*_*I*_ is the activation rate of *Q*_*u*_ playing the role of the integral gain, and *γ*_*c*_ is the dilution rate of the quorum sensing molecules inside the controller cells.

The dynamics of *Q*_*u*_ and *Q*_*x*_ in the population not producing them can be described by

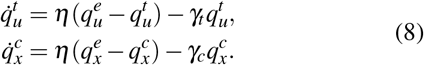

Finally, the dynamics of the quorum sensing molecules in the environment is given by

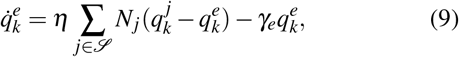

where ∈ {*k x, u*}, 𝒮 = {*t, c*} and *N*_*j*_ is the number of cells in each population. All the other parameters of the consortium have the same meaning as in (1). Note that in the model described in this section, it is assumed to have unlimited cellular resources.

### C. Burden model

Gene expression depends on finite cellular resources, including chemical species, RNA polymerase, and ribosomes [10]. In this work, we focus on the limited availability of free ribosomes – which translate mRNA into amino acid chains to form proteins – as the primary constraint on gene expression. This choice is supported by experimental evidence identifying ribosomes as a key limiting factor in bacterial gene expression during exponential growth [10], [19].

Since ribosomes are finite in number and can only translate one mRNA at a time, their availability directly affects protein production rates throughout the cell. To capture these cross-coupling effects, we employ the *burden* model introduced in [11], which we briefly summarize here for completeness.

Consider the following set of chemical reactions describing the process of synthesis of protein *X*_*p*_

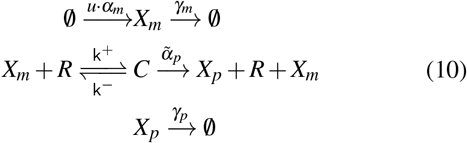

The first reaction is the transcription of the mRNA *X*_*m*_ of the gene *X*, that occurs at rate *α*_*m*_ and depends on the presence of the input signal *u* (e.g. it is constitutively expressed by the promoter or promoted/inhibited by a transcription factor). The mRNA also degrades at rate *γ*_*m*_. The second reaction represents the binding of the mRNA *X*_*m*_ with a free ribosome *R*, forming the translation complex *C*. This reversible reaction happens with forward and reverse rates *k*^+^ and *k*^−^, respectively. Finally, the protein *X*_*p*_ is produced at rate 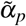 and degrades at rate *γ*_*p*_.

By applying the law of mass action, the dynamics of *X*_*m*_, *C* and *X* can be described by

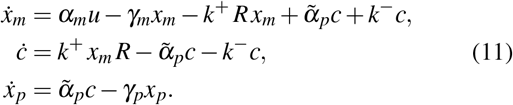

Since the dynamics of the translation complex *C* occur much faster than protein dynamics [20], we can apply a quasi-steady state assumption 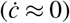, yielding

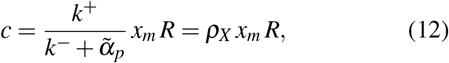

where 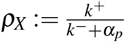 is termed the resource demand coefficient (RDC) associated to protein *X* [11]. Under this assumption, model (11) becomes

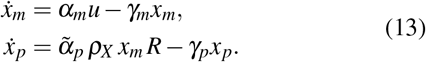

At each time instant a ribosome can be either free or forming the translation complex of a protein. Denoting as *X*_*i*_, *i*∈ {1, …, *n*} all the synthetic proteins produced within the cell, and assuming the total amount of shared ribosomes to be constant and equal to *R*_*T*_, we can write

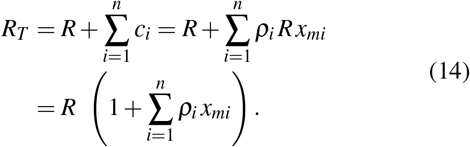

This implies that

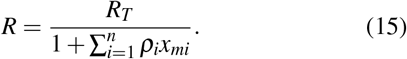

Finally, assuming also the mRNA dynamics to be at quasi-steady state [20], i.e. 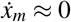, we obtain

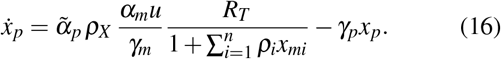

## III. Effects of metabolic burden on the AIC

This section presents mathematical models with burden for both embedded and multicellular AIC, including their steady-state *X*_2_ values. We first review the embedded AIC model with metabolic burden from [11], then derive a resource-aware model for the multicellular implementation.

### A. Embedded AIC with Metabolic Burden

The mathematical model describing the dynamics of the embedded (or single-cell) AIC (1) in the presence of limited resources (i.e. ribosomes) can be obtained by modeling limited ribosomes, as described in equation (16), for all the species implementing the AIC, yielding [11]

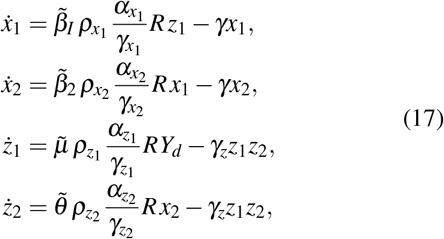

where *α*_*X*_ */γ*_*X*_, with *X*∈ {*X*_1_, *X*_2_, *Z*_1_, *Z*_2_} is the ratio between the elongation and the degradation rate of mRNA associated to gene 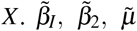 and 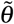 are the protein synthesis rates of *X*_1_, *X*_2_, *Z*_1_ and *Z*_2_, respectively.

The relationship between these parameters and the corresponding ones in (1) can be directly established in the case of abundant resources (i.e. 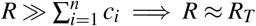

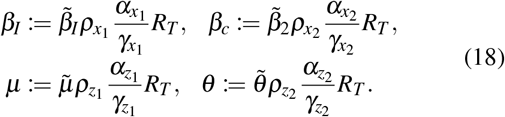

All the other parameters have the same meaning as in (1) and (16).

The expression of *R* in (17) can be obtained from (14) by considering the cascade of genes in the AIC, that is

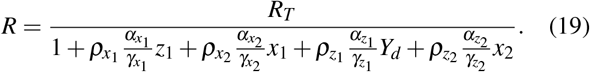

This relationship highlights that the production rate of each protein in (17) depends not only on the concentration of the species that directly activates its expression, but also on the concentration of free ribosomes *R* in (19), which depends on the concentration of the other proteins in the circuits.

Given some fixed value of the reference signal *Y*_*d*_ and provided that the values of the parameters in (17) are such that the system converges to a stable, admissible equilibrium point (i.e. such that *R ≠* 0), the value of the output species *X*_2_ at steady state is given by

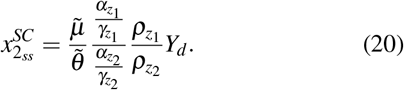

### B. Multicellular AIC with Metabolic Burden

We next derive an aggregate model describing the mean dynamics of species concentrations in the multicellular AIC consortium. We use superscripts *c* and *t* to denote quantities averaged over Controller and Target populations, respectively. The protein concentration dynamics in the Target population are defined by:

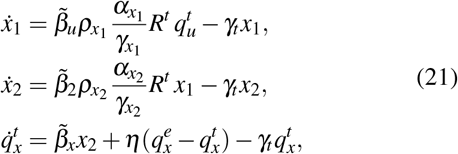

where 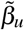 represents the production rate associated to *x*_1_ due to *Q*_*u*_, 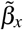 the production rate of *Q*_*x*_ due to *X*_2_, and *R*^*t*^ represents the average concentration of free resources inside the Targets. Quorum sensing molecules are assumed to be synthesized by means of enzymatic reactions, thus they do not utilize ribosomes.

Similarly, the aggregate dynamics in the Controllers is described by

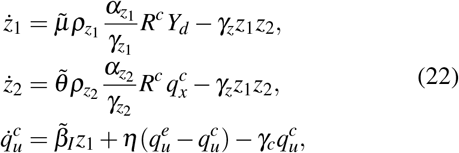

where *R*^*c*^ represents the average concentrations of free resources in the Controllers.

The expressions of *R*^*c*^ and *R*^*t*^ are different from the one defined in (19). Specifically, within the Controllers only *Z*_1_ and *Z*_2_ take up resources, whereas in the Targets *X*_1_ and *X*_2_ are the only species sharing resources.

The relationship that describes the utilization of resources in the *h*-th cell of population *j* ∈ {*c, t*} can be formalized as

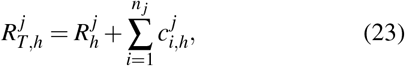

where 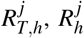, and 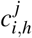 represent the concentration of total resources, of free resources and of translation complexes in population *j*, respectively, and *n* _*j*_ = 2 is the number of chemical species sharing of ribosomes within population *j*.

Averaging (23) over the entire population *j*, e.g. *c*, yields

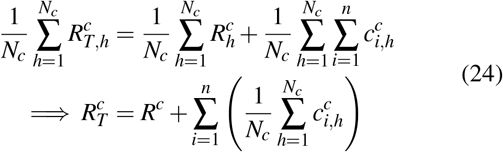

where *N*_*c*_ is the number of cells in the Controller population.

Substituting (12) in (24) in place of the average concentration of translation complex 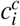 and solving for *R*^*c*^, we get

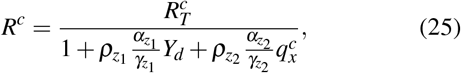

where 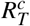 and 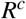 represent the average concentration of total and free resources in the Controller population, respectively.

Following similar steps, we get

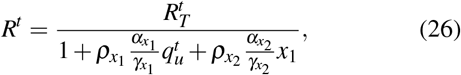

where 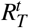 and 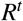 represent the average concentration of total and free resources in the Target population, respectively.

In order to derive the analytical expression of the output at steady state, we make the following assumptions:

**A1**. All cells grow and divide at the same rate. This implies that all cells have the same dilution rate: *γ*_*c*_ = *γ*_*t*_ = *γ*. This assumption holds if cells are of the same biological chassis, and thus they grow and divide at the same rate.

**A2**. The quorum sensing molecules *Q*_*u*_ and *Q*_*x*_ diffuse faster than they degrade. This implies that *η* ≫ Γ_*I*_, with Γ_*I*_:= *γ*_*c*_ + *γ*_*t*_ = 2*γ*, see e.g. [21].

Under assumptions **A1**-**A2** and provided that there exists a stable, admissible equilibrium point, the value of the output species *X*_2_ at steady state is given by

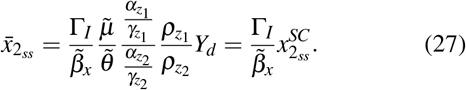

## IV. Robustness analysis and in silico experiments

This section compares embedded and multicellular designs based on their ability to regulate *X*_2_ expression to desired levels under resource constraints. We analyze the structural stability of the desired equilibrium point as total resources *R*_*T*_ vary, and validate both architectures through realistic *in silico* agent-based experiments.

To enable direct comparison, we selected parameters Γ_*I*_ and 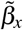 in the multicellular AIC to match the steady-state values 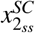 and 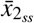 from equations (20) and (27), respectively.

### A. Numerical Bifurcation Analysis

We conducted numerical continuation analysis using MATCONT [22] to identify stable equilibrium ranges for both architectures as total available resources per cell (*R*_*T*_) vary. For the multicellular AIC, we analyzed equations (21), (22), (8), and (9), assuming equal resources in both populations 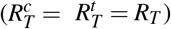. The resulting bifurcation diagram (Fig. 2c) shows a saddle-node bifurcation at *R*_*T*_ = 273.79 nM (marked as LP, limit point) and a Hopf bifurcation at *R*_*T*_ = 1544.76 nM.

**Fig. 2:**
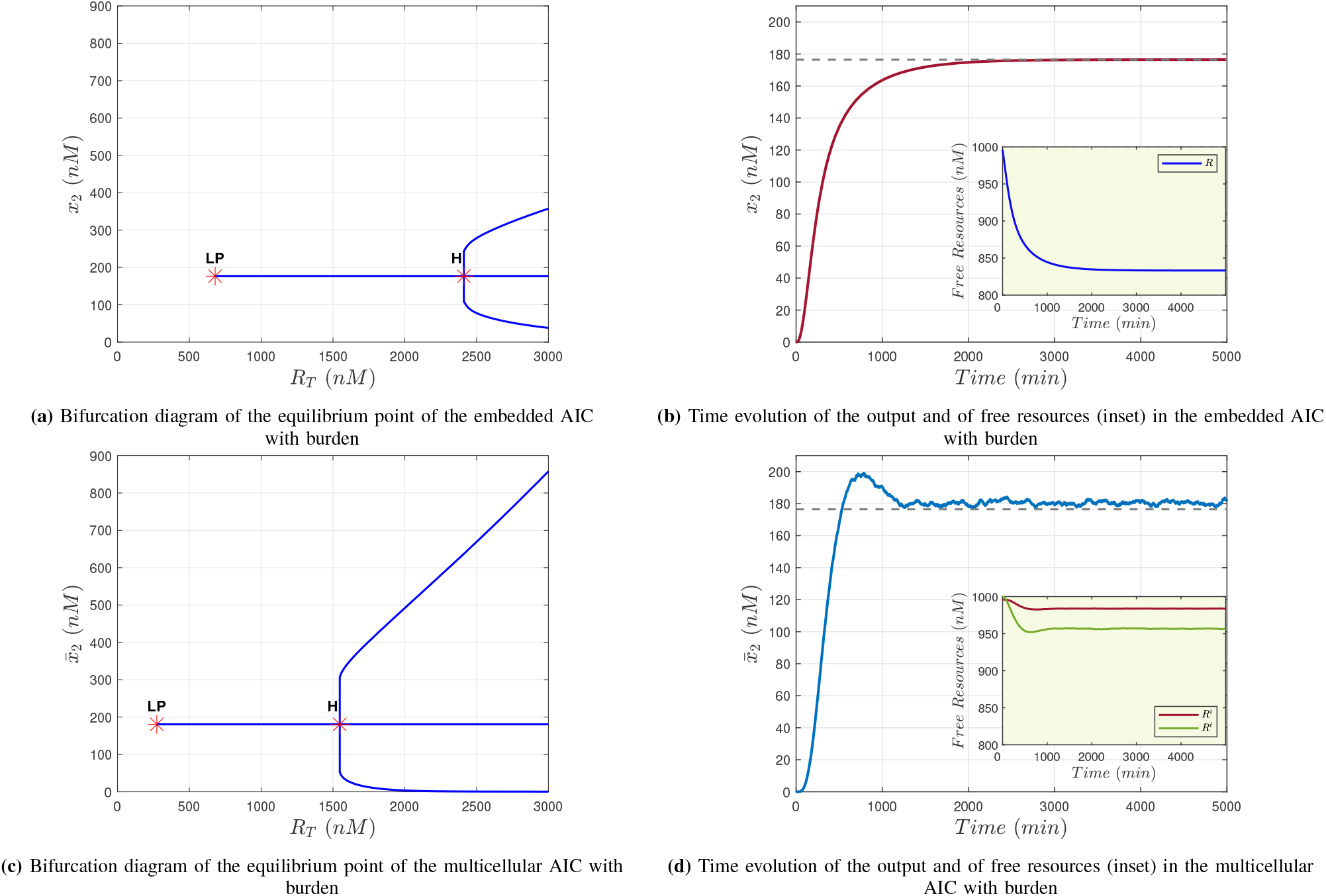
*In silico* experiments on the embedded and multicellular AIC with burden. Bifurcation diagrams of the equilibrium point of the embedded (a) and multicellular (c) AIC when *R*_*T*_ is chosen as the free parameter. The equilibrium points first undergo a Saddle-Node bifurcation at *R*_*T*_ = 679.87 nM and *R*_*T*_ = 273.79 nM, respectively, and then a Hopf bifurcation at *R*_*T*_ = 2411.28 nM and *R*_*T*_ = 1544.76 nM, respectively. Stable equilibrium points exist between the two bifurcations. Time evolution of the output of the embedded (b) and multicellular (d) AIC, respectively. Fluctuations in (d) are due to variation in the number of individuals in each population over time. The introduction of the quorum sensing molecules causes the multicellular steady-state error to be different than zero, in particular *e*_%_ = 2.23%. The value of 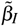 was set to 0.066 min^−1^ in the multicellular scenario, and to 0.0857 min^−1^ in the embedded. Insets in Figs. (b) and (d) show time evolution of the free resources in the embedded and multicellular AIC, respectively. In the multicellular scenario, the *controller* population (red line) consumes 1.23% of initial resources, while the *target* population (green line) consumes 4.34%. In the embedded scenario, consumption settles at 16.3% of initial resources. Simulations via *BSim* were carried out in a chamber of dimensions 20 *µ*m × 15 *µ*m × 1 *µ*m, starting with 20 bacteria (10 for each population in the multicellular scenario) lined up horizontally in the middle of the chamber. Nominal values of the biochemical parameters are reported in Appendix A.

**Fig. 3:**
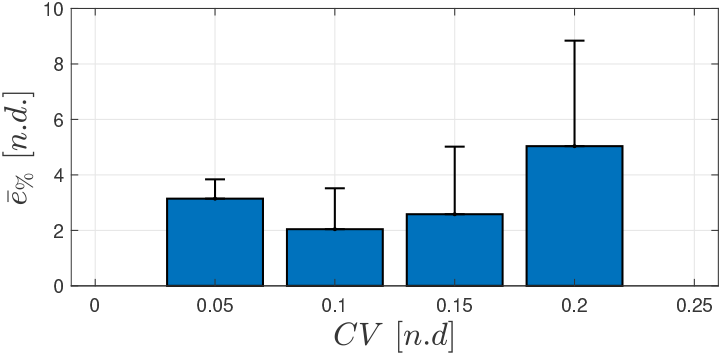
*In silico* robustness experiments in *BSim*. The multicellular architecture’s communication mechanism was tested against cell-to-cell variability. Each parameter, say *α*, was extracted from a Gaussian distribution centered around the nominal value, 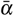, and with standard deviation 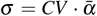, where *CV* represents the coefficient of variation. For each value of *CV* ∈ {0.05, 0.10, 0.15, 0.20}, we evaluated the steady-state percentage error (28) averaged over 10 experiments. The simulations were set up as described in Section IV-B.

The embedded AIC’s bifurcation diagram (Fig. 2a) reveals a stable equilibrium point for *R*_*T*_ ∈ (679.87; 2411.28) nM. While this stability range is 36% wider than the multicellular case, the embedded architecture requires three times more minimum resources to operate. Notably, the multicellular AIC achieves stable equilibrium even under scarcer resources per cell, where the embedded design fails to stabilize. Specifically, the multicellular system operates with a total resource level per cell *R*_*T*_ *>* 273.79 nM while the embedded system requires *R*_*T*_ *>* 679.87 nM. Notice that when self-degradation dynamics of *Z*_1_ and *Z*_2_ are also taken into account in the AIC models (21) and (22), a stable equilibrium point still exists for low *R*_*T*_ values. However, the corresponding steady-state value of *x*_2_ goes to zero as *R*_*T*_ decreases, thus the AIC does no more behave as an integral controller.

### B. In silico experiments

We validated the multicellular architecture using BSim [15], a realistic agent-based simulator for microbial populations that accounts for physical and geometrical effects including cell-environment interactions, cell growth and replication, and chemical species diffusion in the growth medium. Our simulation chamber (20 *µm* × 15 *µm* × 1 *µm*) accommodated approximately 75 cells, with top and bottom openings allowing cell turnover through growth and division. Each simulation began with 20 bacteria arranged horizontally in the chamber’s middle (split equally between populations in the multicellular scenario).

Figures 2d and 2b demonstrate that both architectures successfully regulate *X*_2_ expression to desired levels, though the multicellular implementation shows some fluctuations due to uneven population turnover. However, from the inset plots in Figures 2d and 2b reporting the time evolution of the free resources in the cells, it is apparent that cells in the embedded architecture consume about three times more resources than those in the multicellular one.

### C. Robustness to cell-cell variability

Finally, we tested the robustness of the multicellular architecture’s communication mechanism to cell-cell variability. Specifically, we varied the parameters associated with the production and diffusion of the quorum sensing molecules, such 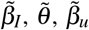, *η*, and the concentration of total resources within the cells, that is, *R*_*T*_. The production rate of *Q*_*x*_ inside the Targets, namely 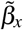, was not affected by perturbations, since it is related to the dilution rate of the chemical species in the cells (see Appendix A). Each parameter, say *α*, was extracted from a Gaussian distribution centered around the nominal value, 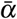, with a standard deviation 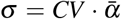, where *CV* represents the coefficient of variation. For each value of *CV* ∈ {0.05, 0.10, 0.15, 0.20}, we performed 10 experiments and evaluated the average steady-state percentage error, defined as

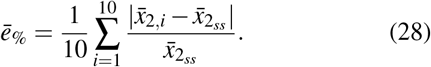

where 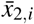 and 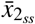 represent the output concentration averaged over the last 200 minutes in the *i*-th experiment and the output at steady-state (see equation (27)), respectively. The simulations show that it guarantees small sensitivity to increasing levels of heterogeneity in the communication mechanism and in the total pool of resources, with the average percentage error never exceeding 10%.

## V. Conclusions

We developed a resource-aware model for a multicellular implementation of the AIC and compared it with its embedded counterpart. Through numerical continuation analysis and realistic *in silico* experiments, we demonstrated that the multicellular AIC maintains stable equilibrium even under scarce resources (low 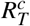 and 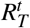), while the embedded implementation requires substantially higher resource levels per cell for operation. This reduction in resource consumption has important practical implications. Lower metabolic burden translates to faster growth rates and improved culture viability, critical factors for industrial applications. Moreover, the reduced resource requirements suggest that multicellular implementations could better accommodate more complex synthetic circuits, where resource competition often leads to circuit failure.

Future work will focus on deriving analytical conditions for parameter selection to guarantee stable equilibrium points. We also plan to extend this analysis to compare the complete multicellular PID controller [14] with its embedded counterpart [16].

## Appendix

### A. Nominal Biochemical Parameters

The nominal biochemical parameters used in Matcont numerical con tinuations and BSim simulations are chosen as: 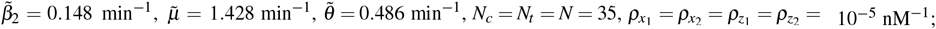 *γ* = 0.023 min^−1^, *γ*_*e*_ = 0.0023 min^−1^, 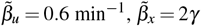, *η* = 2 min^−1^ (taken from [13]); *Y*_*d*_ = 60 nM, *γ*_*z*_ = 0.01 min^−1^nM^−1^ (taken from [16]); *R*_*T*_ = 1000 nM, *α*_*X*_ */γ*_*X*_ = 7, with *X* ∈ {*X*_1_, *X*_2_, *Z*_1_, *Z*_2_} (taken from [10]).

## Notes

This work has received financial support by the European Union - Next Generation EU, under PRIN 2022 PNRR, Project “Control of smart microbial communities for wastewater treatment”.

### Competing Interest Statement

The authors have declared no competing interest.

### Summary of Updates

Modified Section IV with the inclusion of a new paragraph, IV.C, regarding robustness to cell-cell variability. A new figure associated to the added paragraph was included, namely, Figure 3.

